# The RNA-binding protein Puf5 buffers mRNA levels against chromatin-mediated changes in nascent transcription

**DOI:** 10.1101/2020.08.13.249912

**Authors:** David Z. Kochan, Julia S. P. Mawer, Kiril Tishinov, Swati Parekh, Jennifer Massen, Martin Graef, Anne Spang, Peter Tessarz

## Abstract

Gene expression is a dynamic process regulated at all stages, starting with opening of chromatin, transcription, and continuing with mRNA export, translation and, finally, degradation. While there are feedback mechanisms within the system, it is not clear whether these extend to crosstalk between chromatin architecture and mRNA decay. Here, we show that changes in nascent transcription, mediated by mutating H3K56 to alanine, are post-transcriptionally buffered by the Pumilio protein Puf5, which stabilizes transcripts in a context-dependent manner. Depleting Puf5 in an H3K56A background leads to synthetic lethality. This genetic interaction can be explained by a decrease in translation due to downregulation of its direct mRNA targets, largely consisting of ribosomal protein genes. Importantly, we show that this post-transcriptional buffering is not only linked to H3K56A, but may be a more widespread phenomenon that also buffers against an increase in nascent RNA transcription in order to maintain physiological mRNA levels and cellular homeostasis.

## INTRODUCTION

The life of an individual mRNA starts with its synthesis from the DNA template. This process is catalysed by RNA polymerase II and is strongly impacted on by the underlying architecture and dynamics of the chromatin. Indeed, the packaging of DNA into nucleosomes affects all stages of transcription, from transcription factor binding to initiation and elongation (Workman and Kingston 1998). Nucleosomes are subject to a vast array of post-translational modifications (Tessarz and Kouzarides 2014). These regulate (in-)directly the accessibility of the underlying DNA and thus, efficiency of transcription. For instance, acetylation of lysine side chains in histones neutralizes the negative charge of the DNA and weakens the histone-DNA interaction to promote transcription. While we have a solid understanding of the molecular events governing transcription, we still have limited insight into the regulation of mRNA decay. This is largely because standard RNA-seq experiments capture the steady-state levels of RNA and thereby do not shed light on its dynamic lifecycle (Nikopoulou, Parekh, and Tessarz 2019). This shortcoming has led to the development of methodology that combines metabolic labelling of RNAs with genomics to analyse synthesis and degradation rates (Munchel et al. 2011; Miller et al. 2011; Chan et al. 2018). These approaches have helped to reveal the dynamic lifecycle of mRNA, showing that the average mRNA half-live is only a few minutes. This highlights the importance of mRNA degradation pathways in maintaining cellular homeostasis (Chan et al. 2018).

Degradation of mRNA is initiated by the removal of the poly(A) tail, which in yeast is mainly catalysed by the cytoplasmic Ccr4-Not complex (Collart 2016). Subsequently, mRNAs can either be degraded from the 3’ end by the exosome complex or, following mRNA decapping, from the 5’ end by the cytoplasmic exonuclease, Xrn1 (Muhlrad and Parker 1994). Targeting of mRNAs to the Ccr4-Not complex is mediated by several RNA binding proteins, including proteins of the Pumilio family of proteins (Pufs) (Nishanth and Simon 2020). Different Pufs recognize Pumilio-response elements (PREs) of different sizes and have overlapping, but also distinct, target RNAs. Puf5, for instance, can bind around 16% of all yeast mRNAs, a network that is partly shared with Puf3 and Puf4 (Lapointe et al. 2017).

In this study, we set out to address if there is a connection between chromatin dynamics and the complex post-transcriptional network of RNA binding proteins. Could global changes in the underlying histone modification pattern, such as during cancer development or ageing, influence the post-transcriptional life of an mRNA? The rationale for this study comes from observations in mutants affecting acetylation of histone H3K56. Acetylation at this site is important to allow access to chromatin after DNA damage, enhance histone turnover at transcriptionally active chromatin, and to promote nucleosome assembly during S phase (Q. Li et al. 2008; Kaplan et al. 2008; Topal et al. 2019; Wurtele et al. 2012). Despite this important role in shaping chromatin, mutations at H3K56 to alanine or arginine only mildly affect the steady-state transcriptome (Topal et al. 2019; Rege et al. 2015). Intriguingly though, genome-wide nascent transcription is reduced in these mutants (Topal et al. 2019), suggesting that post-transcriptional buffering of mRNA levels must exist. Using a genome-wide genetic screen in *Saccharomyces cerevisiae*, we identify a *puf5* mutant as synthetically lethal with H3K56A, indicating that Puf5 might work as a central player in this buffering system through its targeting of down-regulated nascent transcripts in an H3K56A background. Depletion of Puf5 in this background, leads to the downregulation of ribosomal protein genes, suggesting that the reason for synthetic lethality is a decrease in translation. We further identify mutations in H3K4 that are also genetically linked to Puf5, indicating that the phenomenon of buffering mRNAs upon chromatin-mediated transcriptional change is widespread. Strikingly, in the case of H3K4R, nascent transcription is upregulated, suggesting a context-specific activity of Puf5. This newly discovered post-transcriptional buffering system constitutes another regulatory layer in gene expression to ensure cellular homeostasis.

## RESULTS

### A high-throughput screen identifies synthetic lethality in an H3K56A *puf5Δ* double mutant

The observation that, in an H3K56A yeast mutant, a reduction in nascent transcription does not lead to a reduction in steady-state levels of mRNA, indicates the presence of post-transcriptional buffering of gene expression. To identify proteins that may be involved in such buffering, a high-throughput synthetic genetic array (SGA) screen was performed. The H3K56A mutant was used as a query strain, and screened against the entire yeast knockout (YKO) and Decreased Abundancy by mRNA Perturbation (DAmP) collections (Giaever et al. 2002; Breslow et al. 2008) (Figure 1A). In total, 101 synthetic interactions were identified (Supplementary Table 1).

**Figure 1:**
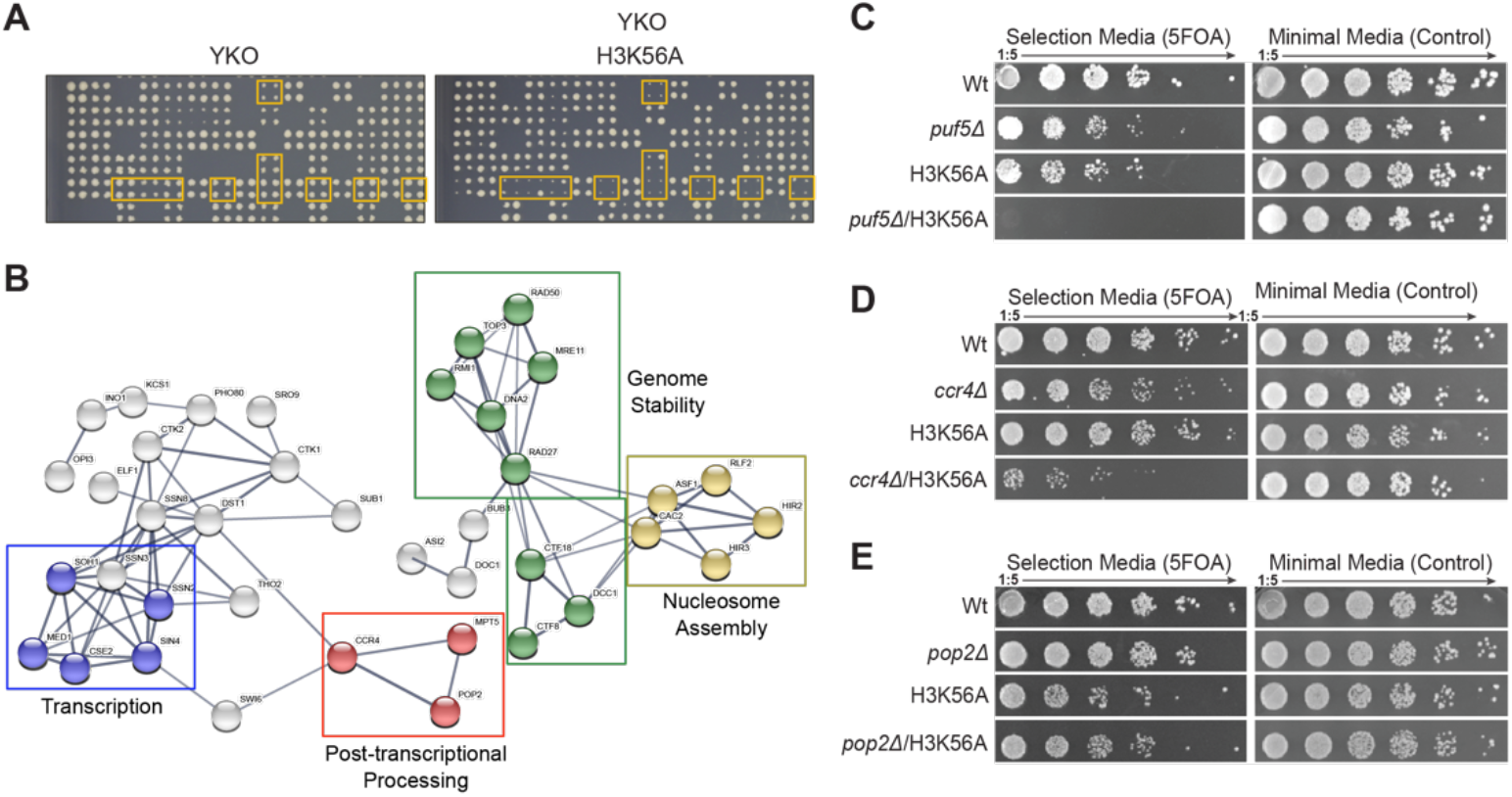
Identification of lethal genetic interaction between *puf5Δ* and H3K56A: A) Exemplary SGA plates highlighting several synthetic genetic interactions between H3K56A and yeast deletion strains (YKO) (right plate). Every mutant is represented in four replicates. B) STRING network representation of protein complexes identified to be genetically linked to H3K56A. GO term for the individual complexes are highlighted by different colours. For details on gene names and GO annotations, please refer to Supplementary Table 1. C-E) Validation of synthetic interactions of the SGA in W303 using a plasmid shuffle system to introduce H3K56A into the indicated genomic deletions of *PUF5* (C), *CCR4* (D) and *POP2* (E).

These included several metabolic complexes, genes required for genome stability, members of the nucleosome assembly machinery, as well as components of the Mediator complex (Figure 1B). Interestingly, constituents of the Ccr4-Not poly-deadenylation complex (Ccr4, Pop2 and Puf5/Mpt5) were also identified (Figure 1B). As the Ccr4-NOT poly-deadenylation complex is known to function in the processing of mRNA, its components could be ideal candidates for a post-transcriptional buffering system of gene expression. To verify the results from the screen, *pop2*, *ccr4*, and *puf5* deletions were generated in a W303 background that allows for the introduction of histone mutations by plasmid shuffle (Tessarz et al. 2014) (Supplementary Table 1). Using this system, the synthetic genetic interaction of H3K56A with *pop2Δ* could not be reproduced (Figure 1C), and only a mild growth defect was observed for H3K56A with *ccr4Δ* (Figure 1D). The H3K56A *puf5Δ* double mutant, however, was confirmed to be synthetically lethal (Figure 1E).

Despite Puf5 being an adapter protein for the core Ccr4-Not machinery, these results suggest that the genetic interaction between H3K56A and Puf5 is largely independent of Ccr4-Not. In support of this, Puf5 has recently been identified as central to regulating mRNA stability in the absence of the Ccr4-Not complex (Wang et al. 2018). Taken together, using a synthetic genetic screen, we have identified a member of the Puf protein family to be synthetically lethal with an alanine substitution in histone H3 at K56. The identification of a gene known to be involved in post-transcriptional control of mRNA levels supports our initial hypothesis that the decreased level of nascent mRNA in an H3K56A mutant might be protected from degradation and thus post-transcriptionally buffered to ensure homeostasis of gene expression and cell survival.

### Cytoplasmic localization and RNA-binding are essential for Puf5 function in an H3K56A background

Several mRNA targets of Puf5 code for proteins involved in the maintenance of chromatin architecture (Wilinski et al. 2015; Lapointe et al. 2017). Therefore, *puf5Δ* could result in changes in chromatin structure that become lethal upon loss of acetylated H3K56 (H3K56ac). To determine whether a *puf5Δ* mutant has aberrant chromatin structure, ChIP-seq was used to assess genome-wide deposition of H3 and H3K56ac. No differences in the genome-wide levels of H3 and H3K56ac were observed in a *puf5Δ* mutant (Figure 2A). There were also no significant changes in H3K56ac levels at transcription start sites (Figure 2A), indicating that *puf5Δ* does not have an impact on overall chromatin architecture. These ChIP-seq results suggest that Puf5 functions downstream of H3K56 and support a post-transcriptional role for this protein.

**Figure 2:**
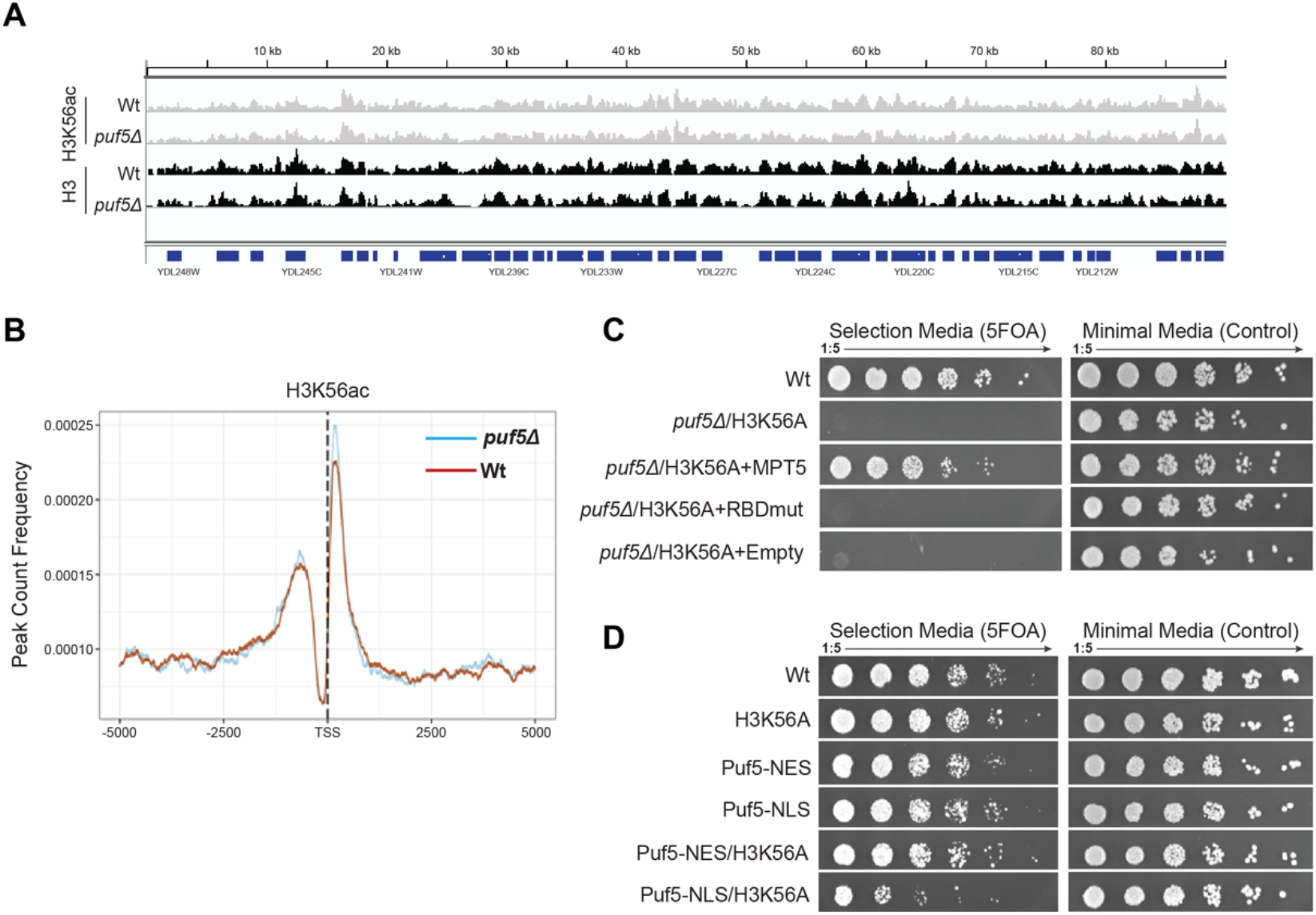
Puf5 function is downstream of chromatin, localised in the cytoplasm and requires RNA binding ability. A) Representative genome browser track of 100kb of chromosome IV showing similar occupancy of H3K56 acetylation and H3 in Wt and *puf5Δ*. B) Metaplot over the TSS +/1 5000bp comparing intensities of H3K56 acetylation in Wt and *puf5Δ* strains. C) Complementation assay using either cytoplasmic (NES) or nuclear (NLS) localised Puf5. D) Complementation assay using a Puf5 mutant unable to bind RNA (S454A, N455A, (Traven et al. 2010)).

Puf5 is an RNA binding protein. As we hypothesise that its function in an H3K56A background is to protect target mRNAs from premature degradation, the RNA binding domain should be essential for this function. To test this, a Puf5 RNA Binding Domain Mutant (RBDmut) (S454A, N455A, (Traven et al. 2010)), was generated and used to complement the H3K56A *puf5Δ* double mutant. As expected, wild-type *PUF5* was able to fully rescue the H3K56A *puf5Δ* lethality, while the Puf5 RBDmut protein was not, confirming that Puf5’s mRNAs-binding ability is essential for H3K56A mutant survival (Figure 2C). Puf5 is localized in the cytoplasm and nucleus. To test if cellular localization plays an important role for the potential buffering of mRNAs, we tagged Puf5 with either a nuclear localization signal (NLS) (Traven et al. 2010) or a nuclear export signal (NES). To evaluate the efficiency of the tag, we additionally added a GFP and monitored Puf5 localization under the microscope, which was consistent with correct targeting of the protein to the respective cellular localisation (Supplementary Figure 1). Interestingly, while Puf5 localized to the cytoplasm complemented a deletion, nuclear Puf5 only partially complemented (Figure 2D), indicating that Puf5 has to be cytoplasmic to buffer the effects of an H3K56A mutation. Together these data suggest that Puf5 functions downstream of H3K56A, in the cytoplasm, and requires its RNA binding ability.

### Rapid degradation of Puf5 in an H3K56A background leads to downregulation of ribosomal protein genes

To identify genes and pathways that rely on the potential buffering effect of Puf5 in the H3K56A background, we used an auxin-induced degradation (AID) system (Tanaka et al. 2015) to quickly degrade Puf5 in the presence of the histone mutation using doxycycline and auxin (Dox/Aux - Supplementary Figure 2A). This system combines transcription shut-off through a Tet-off promoter with auxin-induced degradation that is mediated by an AID tag. However, this system led to a slight overexpression of Puf5. Therefore, we pre-treated cells with low doses of Dox overnight, which decreased Puf5 transcripts, returning them to wildtype levels (Supplementary Figure 2 B). More importantly, in the presence of Dox/Aux, this system recapitulated the synthetic defect between H3K56A and *puf5Δ* in liquid culture and on agar plates (Supplementary Figure 2C, D), while in the absence, the strain showed a similar expression program to an H3K56A strain (Supplementary Figure 2 E)

As additional controls, we performed RNA-seq on wildtype, H3K56A and *puf5Δ* strains. In line with previously published data, we did not observe strong differential gene expression between a wildtype and H3K56A strain (Supplementary Figure 2F), while a *puf5Δ* deletion showed a significant change in the gene expression program (Supplementary Figure 2F). Comparing the individual strains after addition of Dox/Aux, it is obvious that only degradation of Puf5 led to significant changes in the transcriptome, indicating that the combined Dox/Aux treatment only affected the degron strain (Supplementary Figure 2G). Acute depletion of Puf5 in the H3K56A background led to upregulation of 238 genes, while 410 genes were significantly down-regulated (Figure 3A).

**Figure 3:**
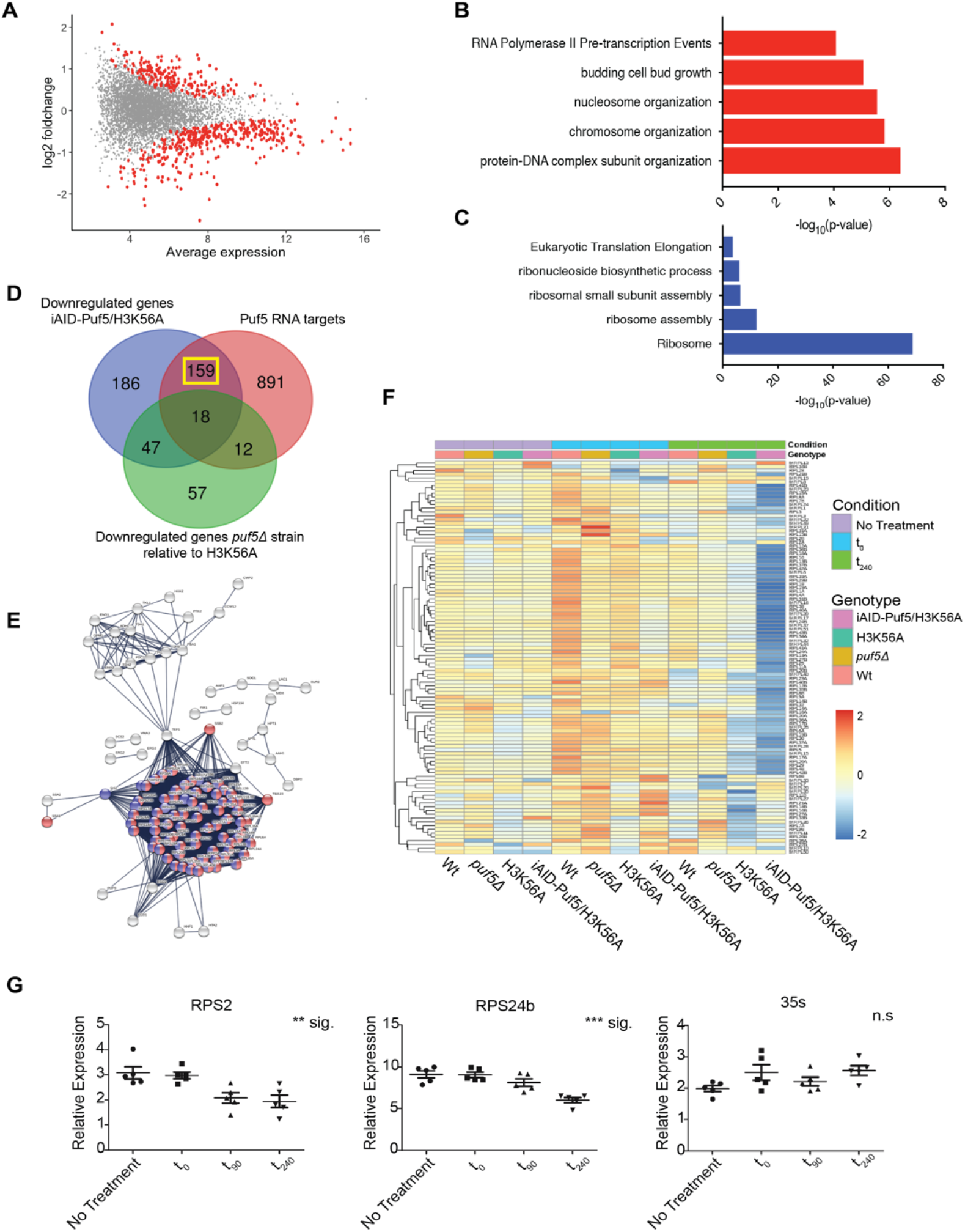
Rapid degradation of Puf5 reveals ribosomal protein genes as targets for Puf5 buffering. A) MA plot comparing H3K56A strains before and after depletion of Puf5. Significantly deregulated genes are highlighted in red. B) Top GO terms for upregulated and C) down-regulated genes upon rapid depletion of Puf5 in an H3K56A background. D) Overlap of down-regulated genes upon Puf5 depletion and its targets, as identified by eCLIP, and differential genes between *puf5Δ* and H3K56A. E) STRING representation of the 159 genes that overlap in (D – yellow box) reveals ribosomal proteins (red/blue) as main, target genes for buffering. For details on the 159 genes highlighted in this network, please refer to Supplementary Figure 2. F) Mean expression level of ribosomal protein genes across the various genetic backgrounds and conditions tested for by RNA-seq. G) RT-qPCR of selected ribosomal protein genes at indicated time points during the depletion of Puf5 in an H3K56A background. 35S rRNA transcription was measured to assess the impact on the nascent transcription of ribosomal DNA. All values are relative to actin. Error bars are SEM, significance was tested using ANOVA (** p < 0.01; *** p < 0.001; n.s. – not significant).

Up-regulated genes were enriched for functions in chromatin organization, cell wall and -cycle related terms (Figure 3B), which might be a potential compensatory response as H3K56A *puf5Δ* cells become very large during the arrest. Strikingly, down-regulated genes were strongly enriched for ribosome-related terms (Figure 3C). To identify genes that depend on Puf5-mediated buffering in the H3K56A background, we intersected all down-regulated genes upon Puf5 depletion in the H3K56A background with i) genes down-regulated between *puf5Δ* and H3K56A and ii) direct Puf5 targets as identified by eCLIP (Wilinski et al. 2015) (Figure 3D). We then focused on the 159 genes that are direct targets of Puf5 and are specifically down-regulated in the double mutant strain (yellow box, Figure 3D), which largely consisted of ribosomal protein genes as well as ribosome assembly and translation initiation factors (Figure 3 E, F).

These results strongly suggest that the synthetic growth defect of Puf5 depletion in the H3K56A background is caused by downregulation of ribosomal protein genes. To assess the specificity of the down-regulation for ribosomal protein genes in the double mutant, we also analysed rDNA transcription. Importantly, while we could confirm down-regulation of ribosomal protein genes, nascent 35S rRNA transcription was not affected by Puf5 depletion (Figure 3G), demonstrating that the down-regulation of ribosomal protein genes is a specific, direct consequence of Puf5 depletion, arguing against an unspecific cellular stress response.

Interestingly, Puf5 shares transcripts of ribosomal protein genes as targets with Puf3 and Puf4 (Lapointe et al. 2017). To test if there were genetic interactions with other Puf proteins, we systematically deleted *puf1* – *puf6* in combination with H3K56A. *puf1*,*2* and *6* deletions did not result in any genetic interaction, but *puf3Δ* and *puf4Δ* showed mild growth phenotypes (Supplementary Figure 3), suggesting a potential partial overlap of substrates with Puf5. Importantly, Puf3/4 only share a fraction of these transcripts with Puf5 (Lapointe et al. 2017), which might explain the much weaker phenotype observed for the deletion of these two Puf proteins.

### Puf5 targets are degraded more slowly in an H3K56A background

To directly test the hypothesis of post-transcriptional buffering of nascent transcripts by Puf5, we asked whether the mRNAs identified as downregulated upon depletion of Puf5 in the H3K56A background are i) de-regulated on the level of nascent transcription and ii) are indeed buffered, i.e. degraded more slowly in the H3K56A mutant in order to maintain a comparable level of these transcripts to wildtype. To address the first point, we made use of a previously published, large dataset comparing nascent transcription in a variety of mutants unable to acetylate H3K56 (Topal et al. 2019) and intersected this dataset with our RNA-seq (Figure 3A). In particular, we used bulk RNA-seq as well as 4sU-seq (nascent RNA-seq) datasets from wildtype and *rtt109Δ* strains, which lack H3K56 acetylation. Indeed, while the steady-state mRNA levels remain stable between wildtype and *rtt109Δ*, the level for nascent transcripts decreases significantly in the histone acetyltransferase deletion strain (Figure 4A). This result strongly reinforces the idea of Puf5-mediated buffering of at least a subset of nascent transcripts. Finally, to directly measure the difference in degradation rates between a wildtype and H3K56A strain, we performed transcriptional shut-off experiments (Collier 2008). Thiolutin was used to block transcription in logarithmically growing yeast and RNA was isolated at 0, 15 and 30 minutes to assess the level of degradation. 18S rRNA served as a reference gene as it was shown to be stable under these conditions (Collier 2008). In the absence of thiolutin, transcripts were stable (Supplementary Figure 4). In line with the hypothesis of post-transcriptional buffering, H3K56A yeast showed slower degradation rates of ribosomal protein genes (*RPL14A*, *RPS9B*, *RPL4B*). SCR1, which was one of the top down-regulated genes did not show any difference, which suggests that it is not buffered, but represents a secondary effect of the Puf5 depletion. On the other hand, *SCR1* has a very long half-life and we might not be able to measure its degradation using transcription shutoff experiments. Importantly, genes that were not impacted by Puf5 depletion in an H3K56A background did not show any differences in degradation rates (*Ubc6*, *Actin*).

**Figure 4:**
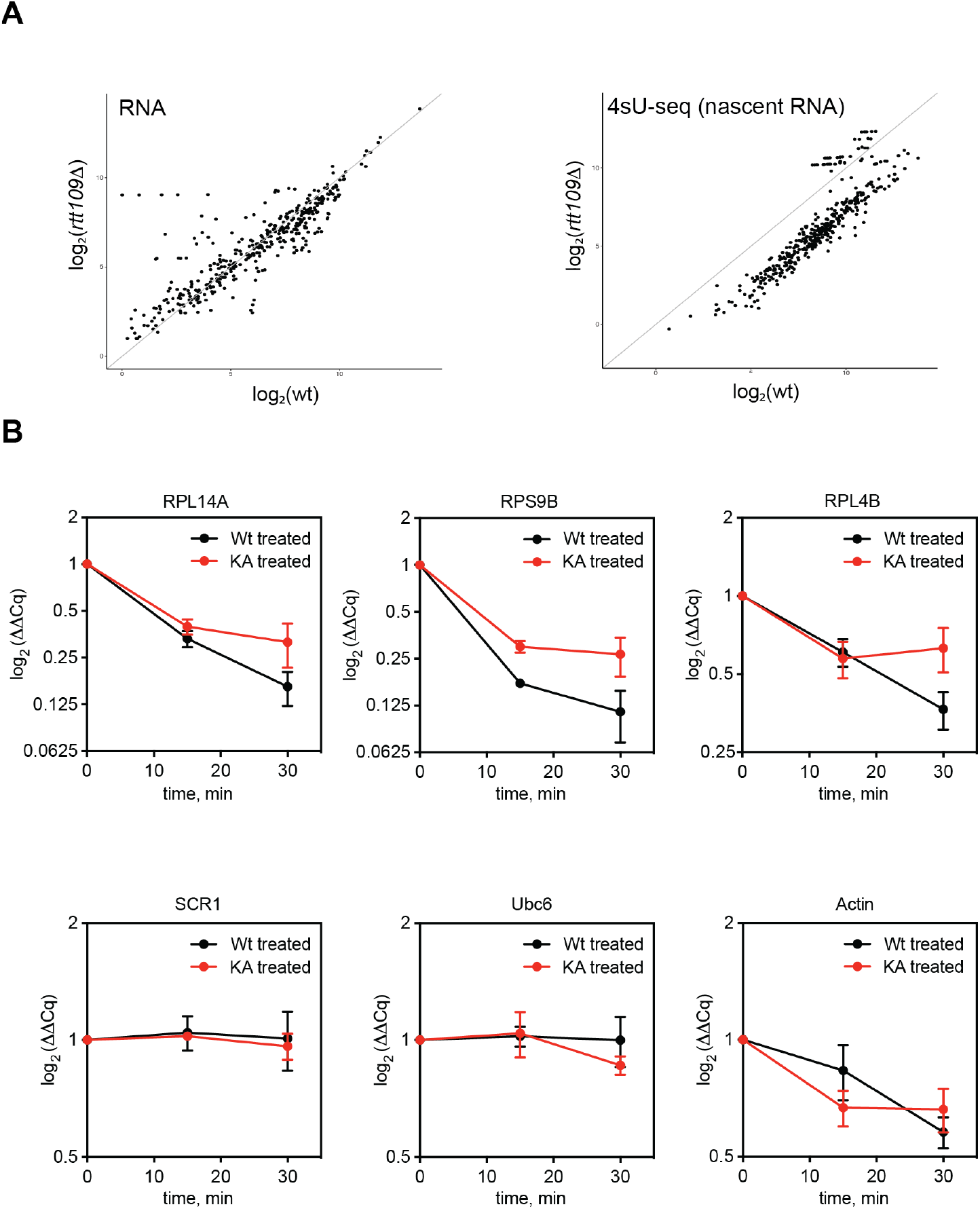
Puf5-buffered mRNAs are degraded more slowly in an H3K56A background. A) Comparison between bulk (steady-state) mRNA levels in Wt and *rtt109Δ* mutants with nascent RNA (4sU-seq) levels in the same strains. Plotted are mRNAs identified to be down-regulated upon Puf5 depletion in an H3K56A background (see Figure 3). Data was taken from Topal *et al*, 2019. B) Transcription shutoff experiments. Cells were grown to mid-log phase before transcription was halted by addition of Thiolutin. RNA was extracted at the indicated time points and RT-qPCR was performed. Gene names are given above the graph. Data is shown relative to 18S and time point t=0. N=5 and error bars are SEM.

### Post-transcriptional buffering by Puf5 is more widespread and also protects against upregulated nascent transcription

Another interesting aspect to explore was how widespread this phenomenon of mRNA buffering is. To start to address this question, we performed a mini-screen with histone mutations that change amino acids known to be involved in either DNA replication (H4K5,12) or transcription (H3 K4, 16, 36 and 79). Of these mutations, H3K4R showed a strong synthetic phenotype in combination with *puf5Δ* (Figure 5A and Supplementary Figure 5A-D) that was similar to the one observed with H3K56A, thus indicating that the Puf5 buffering effect might be more widespread. No genetic interaction with *ccr4Δ* or *pop2Δ* was observed (Figure 5B, C), indicating that the role of Puf5 is independent of its classical function as an adaptor for the Ccr4-Not complex – similar to the role in the H3K56A background. However, we observed two very obvious differences compared to the H3K56A *puf5Δ* genetic interaction: First, the cytoplasmic localisation of Puf5 is not sufficient to complement an H3K4R/*puf5Δ* double mutant (Figure 5D).

**Figure 5:**
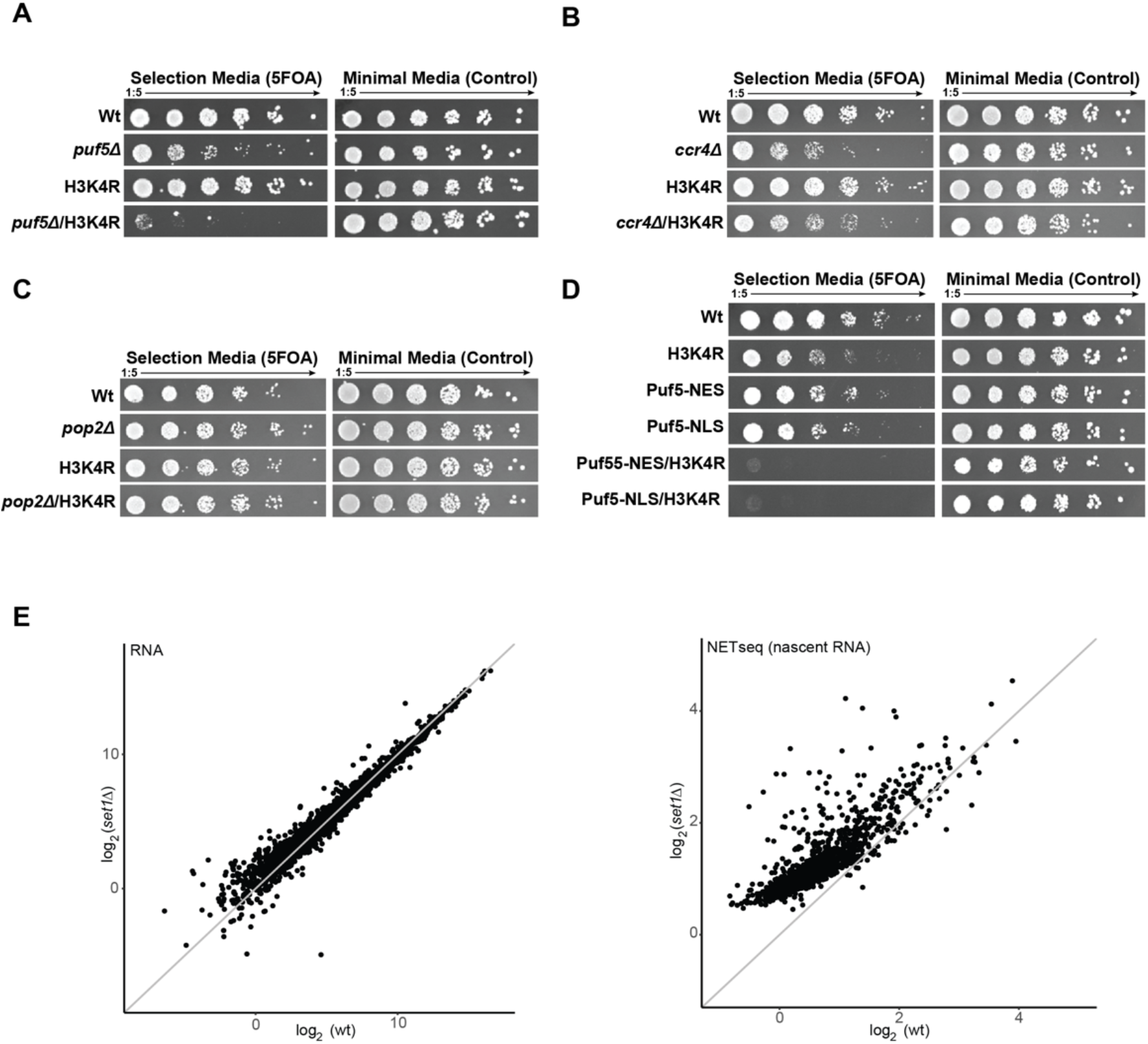
Deletion of puf5 is synthetically lethal with H3K4R. A) Spot test showing synthetic lethality between H3K4R and *puf5Δ*. No synthetic genetic interaction was seen by combining an H3K4R mutant with deletions in B) *ccr4* and C) *pop2*. D) Complementation assay to test for a sufficiency of Puf5 being recruited to cytoplasm (NES) or nucleus (NLS). E) Comparison of RNA steady-state and nascent RNA levels between deletion of the H3K4 methyltransferase, Set1, and wildtype cells.

Second, the targets of Puf5 in an H3K4R background are not transcripts that stem from downregulated nascent transcription. We compared bulk RNA-seq and nascent RNA (NET-seq) data (Churchman and Weissman 2011) for a deletion of SET1, the sole methyltransferase in yeast responsible for methylating H3K4 (Santos-Rosa et al. 2002; Briggs et al. 2001), thus mimicking an H3K4R mutation. In *set1Δ*, nascent RNA was upregulated compared to wildtype levels, whereas steady-state mRNA levels were largely indistinguishable from wildtype cells (Figure 5 E, F), suggesting that Puf5 makes context-specific decisions with respect to degradation or stabilisation of mRNA.

## DISCUSSION

Using a genome-wide genetic screen, we uncovered a post-transcriptional buffering system that can counter defects in nascent transcription to maintain steady-state level of mRNAs. This is the first report that connects chromatin mediated changes to nascent transcription with a post-transcriptional surveillance system. This work adds to previous observations that Puf proteins can make context-specific decisions to degrade or stabilize transcripts in different environmental conditions (Lee and Tu 2015; Wang et al. 2018). Future work will have to address how Puf proteins are able to make these complex decisions.

The two mutations found to be synthetically lethal with *puf5Δ* are H3K4 and H3K56. Interestingly, both residues and their modification have recently been implicated in playing a major role in buffering gene expression during DNA replication, in which the transcript number of a given gene remains the same although the underlying DNA sequence is duplicated (Voichek et al. 2018; Voichek, Bar-Ziv, and Barkai 2016). It is tempting to speculate that Puf5 might be involved in this process too. Unfortunately, it is difficult to evaluate this in the context provided here, as most genes detected to be downregulated upon Puf5 depletion in the H3K56A background, were originally excluded from the cell-cycle buffered transcripts since ribosomal protein genes strongly change in abundance during the cell cycle (Voichek, Bar-Ziv, and Barkai 2016).

In addition to this role in transcriptional homeostasis during the yeast cell cycle, a post-transcriptional buffering system would constitute an important system to maintain transcriptional stability under conditions in which transcriptional rates are impacted, e.g. upon exposure to UV irradiation (Andrade-Lima *et al*, 2015). Also, during ageing, many post-translational modifications on histones become deregulated and change in abundance. The levels of H3K56ac, for instance, are already strongly decreased in the juvenile age of yeast after around 6-7 divisions (Dang et al. 2009). A buffering system might be able to compensate for this loss of histone acetylation. This speculation is supported by the following observations: i) over-expression of Puf5 prolongs replicative lifespan (Kennedy et al. 1995; Kaeberlein and Guarente 2002; Kaeberlein et al. 2004), ii) while deletion of Puf5 strongly decreases replicative lifespan (Kaeberlein and Kennedy 2005).

The family of Pumilio RNA-binding proteins is highly conserved among most eukaryotes (Wang et al, 2018). It will be exciting to see if this phenomenon of post-transcriptional buffering extends not only to other chromatin-mediated effects and cellular states, but also to other organisms. The increased availability of techniques to measure nascent transcription, will allow us to identify conditions and settings that impact nascent transcription based on environmental cues, which are subsequently buffered to sustain the mRNA pool and maintain cellular homeostasis.

## MATERIALS AND METHODS

### Strains and Plasmids and reagents

Genotypes of strains and yeast plasmids used in this work are listed in Supplementary Table 3 and 4. All chemicals used in this study were purchased analytical grade from either Sigma Aldrich or Carl Roth except for the following: Drop Out Mix for yeast synthetic medium was from US Biological Life Sciences (D9543-01). 5-FOA was bought from Cayman Chemical (17318). Ampure XP beads were from Beckman Coulter (A63881), Protein-G coupled dynabeads from Thermo Fisher (10009D) and Thiolutin from Abcam (ab143556). Secondary antibodies against rabbit (7074S) and mouse (7076S) IgG coupled to HRP were purchased from Cell Signaling. Antibodies against H3K56ac were from Active Motif (39281), G6PDH (A9521) and c-Myc (MABE282) from Sigma, iAID-tag (M214-3) from MBL International and β-actin (GTX109639) from GeneTex.

### Yeast protocols

If not stated otherwise, all strains used were derivatives of W303. Integrations and deletions were performed using one-step PCR-based methods (Janke et al. 2004; Longtine et al. 1998; Tanaka et al. 2015). Yeast were grown in YPD medium. Protein was extracted from yeast using sodium hydroxide lysis and TCA precipitation as previously described (Knop et al. 1999). RNA was extracted using the hot phenol approach (Schmitt, Brown, and Trumpower 1990). For spot tests, cells were grown over-night, diluted to an OD_595_=1 and 5-fold serially diluted.

### Auxin-mediated degradation of Puf5

Single yeast colonies were picked and inoculated overnight in 5 mL of YPD media at 30°C. The next day, cultures were diluted in 20 mL of YPD media to an OD_595_ of ~0.15 and grown at 30°C until an OD_595_ of 0.6-0.8. From each cell culture, 10 mL were collected for RNA extraction and 1 mL of cell culture was diluted in 4 mL of YPD media containing 0.25 μg/mL of Dox and incubated overnight at 30°C. The following day, the overnight cultures were diluted in 40 mL of YPD media containing 0.25 μg/mL of Dox to an OD_595_ of 0.2 and incubated at 30°C to an OD_595_ of 0.6-0.8. From each cell culture, 10 mL were then collected for RNA extraction and the remaining cell culture was diluted in 40 mL of YPD media containing 40 μg/mL of Dox and 1mM Aux to an OD_595_ of ~0.2. The cell cultures were then further incubated at 30°C and additional samples were taken at 90 and 240 min.

### 3’-RNA sequencing and analysis

Bulk 3’ RNA-seq libraries were prepared following the SCRB-seq protocol (Bagnoli et al. 2018). A detailed protocol is available on request from the corresponding author. Briefly, 50 ng per sample was reversed transcribed using a polyT_30_ oligo containing a 8bp barcode for later identification and a unique molecular identifier (UMI). RT used the template switching. After the RT step, all reverse transcribed samples were pooled into a single 2 mL Eppendorf tube and DNA was purified with Ampure XP (supplier, ordering info) beads at 1:1 vol/vol. Purified DNA was eluted in 17 μL H_2_O and treated with exonuclease (New England Biolabs, ordering info missing). Subsequently, cDNA was amplified and the final PCR product was purified with Ampure XP beads at a volume to volume ratio of 0.8. Finally, 0.8ng cDNA was tagmented in five replicates. Quality of library was assessed on an Agilent Tapestation and sequenced on an Illumina HiSeq4000 in a 2×75bp mode to around 1 million fragments per sample. The sequenced reads were processed using zUMIs (version 2.0.6) (Parekh et al. 2018) with STAR (version 2.6.1a) (Dobin et al. 2013), samtools (version 1.9) (H. Li et al. 2009) and featureCounts (Liao, Smyth, and Shi 2014) from Rsubread (version 1.32.4) . The reads were mapped to the yeast genome (R64) with the Ensembl annotation version 91. The genes were filtered using the "filterByExpr" function of edgeR (Robinson, McCarthy, and Smyth 2010) with the min.count=5. The differential gene expression analysis was carried out using limma-voom (Ritchie et al. 2015; Law et al. 2014) approach at the adjusted p-value of 0.05. Obtained sets of differentially expressed genes were further analysed, e.g. through gene ontology (GO) enrichment analysis. Differential gene expression analysis for all conditions can be found in Supplementary Table 5. GO term enrichment was performed using Metascape (Zhou et al. 2019) and interaction networks were generated using STRING (Szklarczyk et al. 2019).

### ChIP-sequencing and analysis

Chromatin immunoprecipitation and library production was performed as described (Tessarz et al. 2014). The fastq reads were mapped to yeast genome (R64) using bowtie2 (Langmead and Salzberg 2012) local alignment and duplicates were then removed using MarkDuplicates program of Picard Tools. The peaks were called using MACS2(version 2.1.1.20160309) (Gaspar 2018), with settings –nomodel, --extsize 150, -B -q 0.05 --keep-dup 1. The peaks were annotated using the ChIPseeker package(Yu, Wang, and He 2015). The differential analysis between Wt and *puf5D* was carried out using edgeR(Robinson, McCarthy, and Smyth 2010). The normalising factors were calculated using “RLE” method within “calcNormFactors”, tagwise dispersion trend was estimated using the default parameters in “estimateDisp” function and a generalised linear model was then fit on the data using “glmQLFit” function in robust mode. The genome binning for differential analysis was done using csaw(Lun and Smyth 2016) with the binsize of 1000 and global window filtering with the minimum enrichment of 2 over background.

### Transcriptional Shutoff

Cells were grown in 100 ml of YPD (30°C and 200 rpm) to mid-log phase (OD_600nm_ 0.5-0.6). Cultures were then split and either DMSO or 3 ug/ml of thiolutin (Abcam ab143556, reconstituted in DMSO) were added. Cells were sampled at time-points 0, 15, 30, and 60 minutes. Total RNA was extracted using the hot phenol method (Schmitt, Brown, and Trumpower 1990). RT was performed using random hexamers with 0.5-1 ug of total RNA. qPCR was performed and the delta delta C_t_ method (ref missing) was used to measure relative expression. 18S rRNA and time-point 0 were used as reference.

### Datasets and Availability

ChIP- and RNA-seq data generated in this study is available at ArrayExpress: XXXX. Bulk and nascent RNA (4sU) sequencing data from wild type and *rtt109Δ* mutants were downloaded from GSE125843 (Topal et al. 2019). Bulk RNA-seq for wildtype and *set1Δ* was from GSE52086 (Martin *et al*, 2014) and NET-seq from GSE25107 (Churchman and Weissman, 2011).

## ACKNOWLEDGEMENTS

We would like to thank all members of the Tessarz lab for discussion. Sequencing was performed at the Genomics Core Facility of the Max Planck Institute for Molecular Genetics (Berlin, Germany). Imaging was performed in the FACS & Imaging Core Facility at the Max Planck Institute for Biology of Ageing. This work was funded by the Max Planck Society (to P.T.) and the DFG (TE1079/2-01 – to P.T.).

## AUTHOR CONTRIBUTION

Conceptualisation: D.Z.K., J.S.P.M. and P.T.; Methodology: D.K., J.S.P.M. and M.G.; Investigation: D.K., J.S.P.M., K.T., J.M.; Formal Analysis: S.P., J.M.; Supervision: A.S. and P.T.; Project Administration and Funding Acquisition: P.T.; Writing of Manuscript: J.S.P.M. and P.T.

---

**Supplementary Figure 1:**
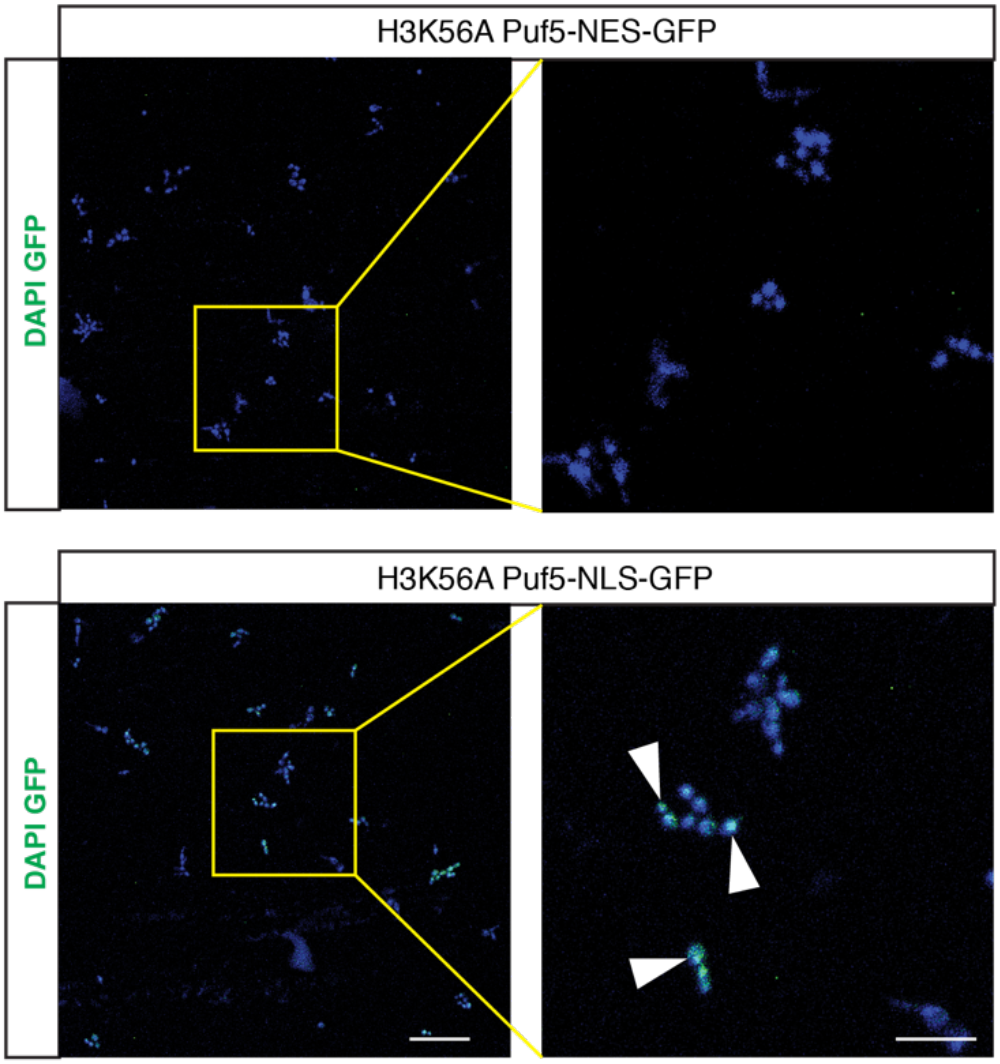
Subcellular localisation of Puf5 fused to NES or NLS, respectively. DAPI served as counterstain for the nucleus.

**Supplementary Figure 2:**
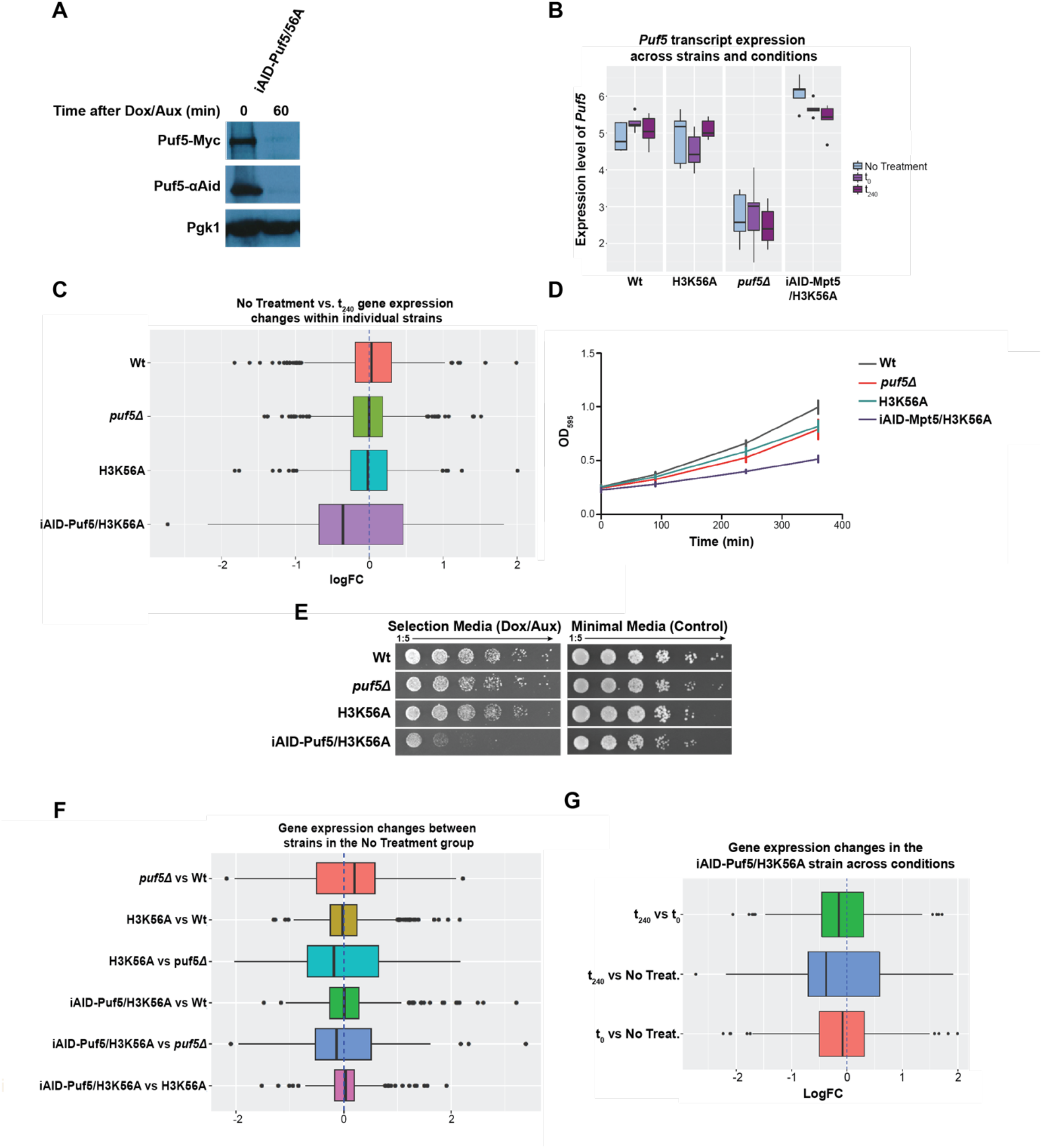
A) Western blot against N-terminal 3xAID-tag and C-terminal myc-tagged Puf5 before and 60min after 40μg/ml and 1mM auxin addition. B) Expression level of Puf5 in the individual strains used in the RNA-seq upon Puf5 depletion, with or without Dox/Aux addition. C) Differential gene expression changes comparing the individual strains used in the RNA-seq experiment to assess downregulation upon rapid Puf5 depletion. Impact of Dox/Aux addition on the growth of yeast in liquid (D) and solid medium (E). (F) Differential gene expression changes upon treatment of Dox/Aux in the individual strains. (G) Differential gene expression changes upon Puf5 depletion in an H3K56A background.

**Supplementary Figure 3:**
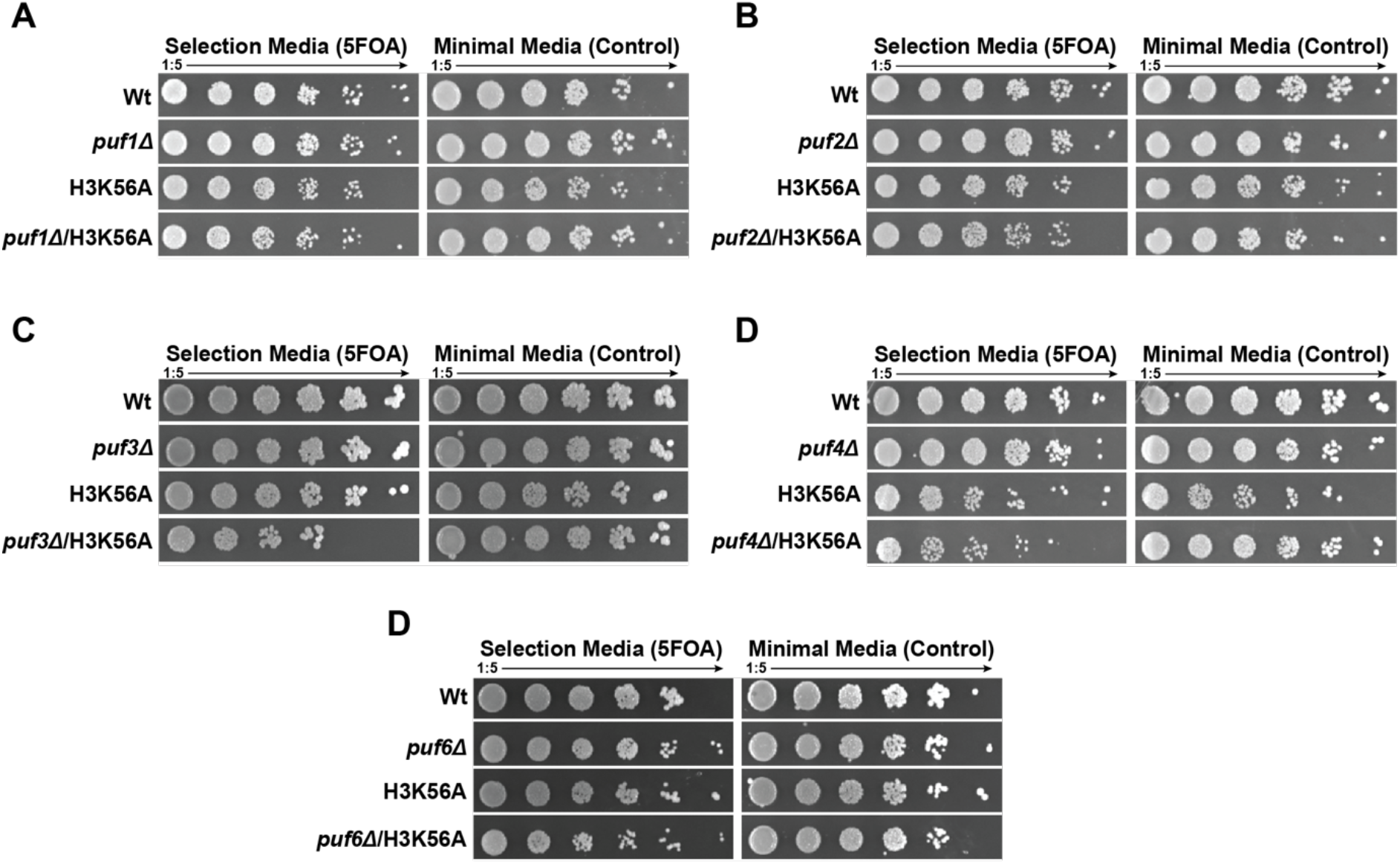
H3K56A shows very mild genetic interactions with PUF3 and PUF4. Spot tests using the histone shuffle strain in combination with A) *puf1Δ*, B) *puf2Δ*, C) *puf3Δ*, D) *puf4Δ* and E) *puf6Δ* deletions.

**Supplementary Figure 4:**
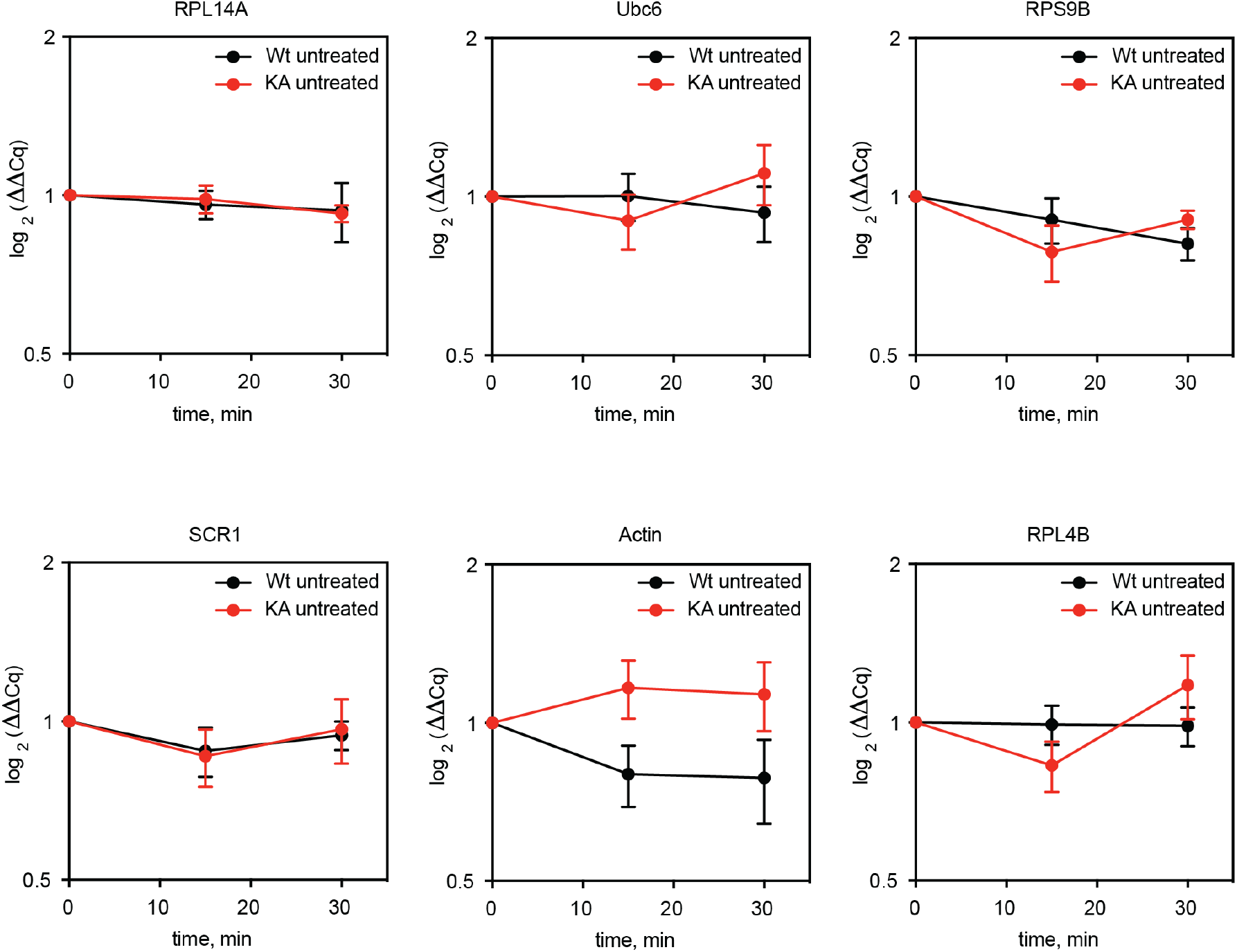
Untreated controls for thiolutin-mediated transcription shutoff (Figure 4). RNA was extracted at the indicated time points and RT-qPCR was performed. Gene names are given above the graph. Data is shown relative to 18S and time point t=0. N=5 and error bars are SEM.

**Supplementary Figure 5:**
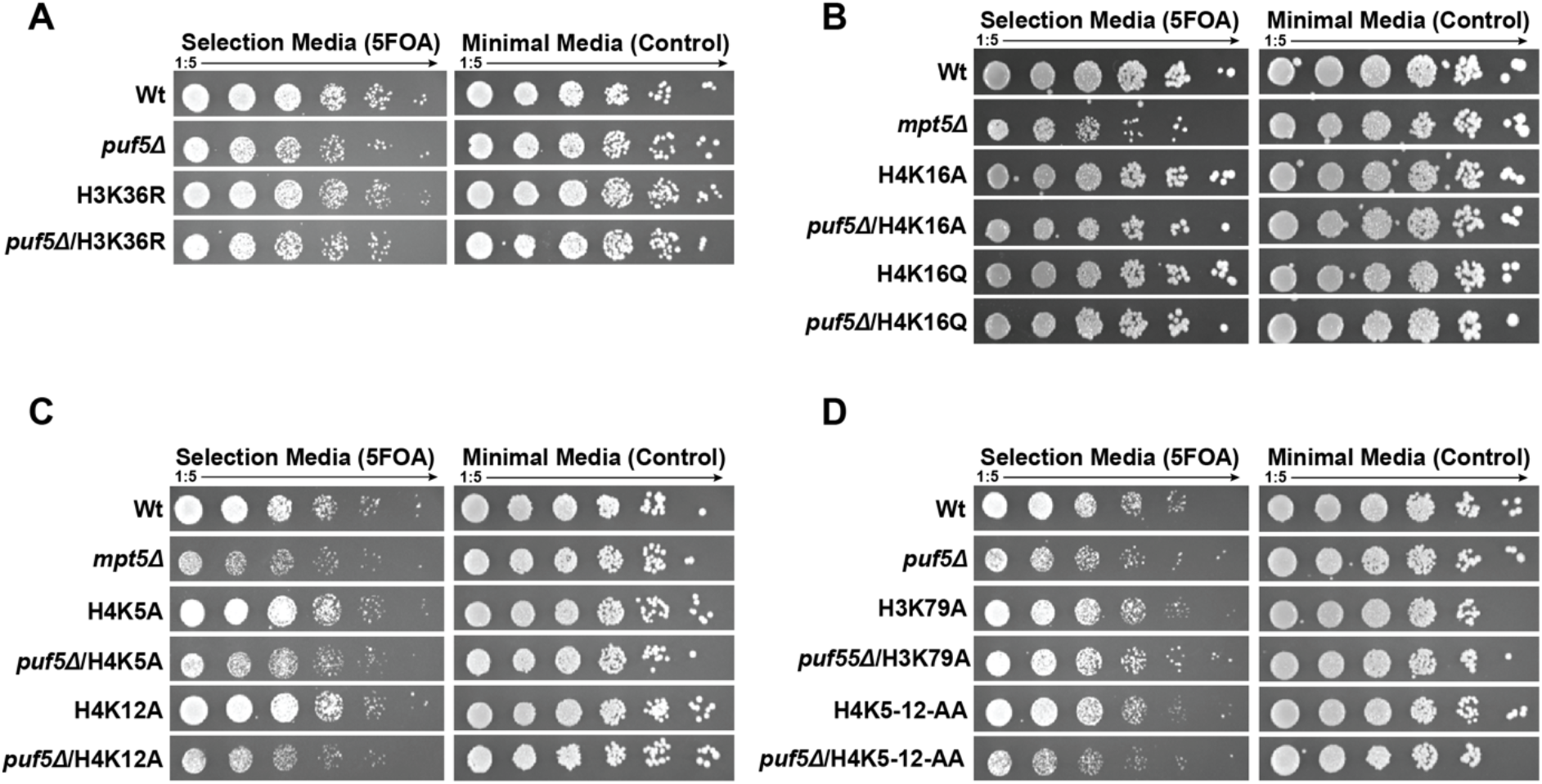
Puf5 is synthetically lethal with an H3K4R mutation. A-E) Various histone mutations of sites involved in transcription and replication were shuffled into the *puf5* deletion strain. F) To complement a *puf5Δ/*3K4R strain, Puf5 has to be present in the cytoplasm as well as nucleus, indicating a different mechanism to the H3K56A mutation.

